# A NOVEL METHOD FOR MEASURING TEXTURE USING A FOOD PROCESSOR

**DOI:** 10.1101/060731

**Authors:** Jiro Kanamori

## Abstract

Measurement of food texture is becoming increasingly important to help prevent mis-swallowing in elderly individuals. However, it is difficult to estimate food texture for mastication and swallowing. In this study, a mixing-recording method was applied to measure food texture. A commercial food processor was used to homogenize food gels, and the torque was monitored as an electric current value, which was correlated with the viscosity of Newtonian fluid. Agar and gelatin gels at several concentrations were applied to the mixing-recording system, and individual mixing curves were obtained. The current values had a different tendency at each mixing stage. At the beginning of the mixing, when lumps of the original gel were present, current values of gelatin were higher than those of agar. From the middle of mixing, the fractured pastes of high-concentration agar gels had little mobility, and current values decreased, indicating that gelatin was more cohesive than agar. The current values of low-concentration agar and gelatin were correlated with the hardness of the gels and were aligned on the same correlation curve, indicating that the viscosity of the fractured paste was correlated with the original gel hardness under certain conditions. Moreover, measurements of agar gels of different sizes revealed that the texture was size dependent. Thus, the mixing-recording method had several advantages over conventional methods, including time- and size-dependent fracturing and bolus properties, and could be useful for the measurement of food texture in terms of mastication and swallowing.

## PRACTICAL APPLICATIONS

The author established a novel method for measuring food texture. The mixing-recording method revealed the physical properties of gels during the fracturing process. The results revealed both similarities and differences between agar and gelatin gels in a different way than conventional methods. Thus, this method could be useful for measurement of food texture to predict mastication and swallowing properties.

## INTRODUCTION

Food is chewed and swallowed as a bolus, the physical properties of which influence the risk of mis-swallowing and aspiration pneumonia (Groher, 1987). The relationship between food properties and mis-swallowing is one of the most important problems associated with food intake in elderly individuals. Mis-swallowing can be induced by various food properties, e.g., adhesiveness, such as that of rice cakes, and noncohesiveness, such as that observed for water and flour (Kumagai and Tanigome, 2011).

Many methods have been established to measure the physical properties of food associated with mastication and swallowing. Viscosity measurement at a shear rate of 50 (1/s) for liquid food is recommended by the American Dietetic Association (2002) and the Japanese Society of Dysphagia Rehabilitation (2013); however, these measurements do not always correlate with human sensory data (Yamagata and Kayashita, 2015). Although many researchers have used the corn plate viscometer, this instrument is difficult to apply to heterogeneous food with solids because of the narrow gap. Texture profiling analysis (TPA), measurement by multiple compressions to calculate several parameters, including hardness, cohesiveness, and adhesiveness, is often used for solid foods (Friedman et al., 1963) and can be used for the classification of hospital foods empirically good for consumption in patients with dysphagia (Sakai et al., 2006). The Japanese Ministry of Health, Labour, and Welfare (2009) adopted TPA as the standard for analysis of foods for patients with dysphagia. However, although TPA has yielded good results for the development and monitoring of foods for patients with dysphagia at hospitals, there are several challenges associated with TPA (Nishinari et al., 2013). For example, the hardness values for some types of foods, such as rubber-like gels and clay-like foods, do not indicate the breaking force, which is related to a sense of hardness. When TPA is applied to liquid foods, the value of cohesiveness is too high, despite their noncohesiveness. Other measurements based on human mastication and swallowing, i.e., sensory evaluation, videofluoroscopy, measuring muscle potential, flow velocity by pulse Doppler analysis, etc., have also been used (Chen, 2009). Although data from these methods may seem to be reliable because they are derived from real human mastication and swallowing, there are many problems with these data, such as high cost, poor accuracy, and individual differences. More reliable, accurate, and simpler methods are needed to measure food texture in relation to mastication and swallowing.

The mixing-recording method (measuring physical properties by monitoring of the mixer torque during mixing) has been used for a long time to measure wheat dough (Bloksma and Bushuk. 1988). Moreover, nonwheat food dough can be analyzed using a food processor as a mixer (Kanamori, 2016). Food processors (i.e., cutting mixers or food choppers) are kitchen appliances with cutting edges that rotate at a high speed. Because the fracturing of solid food can be monitored by the mixing-recording method, I hypothesized that this method could be used to obtain information regarding mastication and swallowing properties.

Agar and gelatin are often used to study the swallowing properties of foods. Gelatin is known to be more cohesive than agar; however, there are no differences in the flow velocity of a bolus, as measured by the pulse Doppler method (Moritaka and Nakazawa, 2010).

In this study, a mixing-recording measurement method was established for application to agar and gelatin gels of several concentrations and sizes. The applicability of the method for food texture analysis was also evaluated.

## MATERIALS AND METHODS

### Materials

Agar and gelatin were high-grade reagents purchased from Kishida Chemicals Co. Ltd. (Tsukuba, Japan). Glycerin was a special-grade reagent purchased from Wako Chemicals. Co. Ltd. (Tokyo, Japan). Silicone oil of low viscosity (52 mPa·s; KF-98-50CS) and high viscosity (3300 mPa·s; KF-98-3000CS) was obtained from Shin-Etsu Chemical Co., Ltd. (Tokyo, Japan).

### Texture measurement

Sample preparation for texture measurement was carried out according to the methods of Moritaka and Nakazawa (2010) with minor modification. The agar suspension was heated at 105°C for 10 min by autoclaving, and the gelatin suspension was heated at 60°C for 30 min in a water bath. The solutions were placed in an acrylic container 40 mm in diameter and 15 mm in height and stood for 15 h at 10°C. Texture measurement for the TPA method was performed according to the methods of the Ministry of Health, Labour, and Welfare of Japan (2009), except that compression strains of 67% and 90% of sample height were used. A creep meter (RE-3305; Yamaden Co. Ltd., Tokyo, Japan) was used at a rate of 10 mm/s with an acrylic plunger (20 mm in diameter and 8 mm in height). Measurements were repeated five times.

### Mixing-recording system

A mixing-recording system was used as described previously with the following modifications (Kanamori, 2016). The system consisted of an electric power source (UPS310HS; Yutaka Electric Mfg. Co. Ltd., Tokyo, Japan), a power transformer (Boltslider N-130-10; Yamabishidenki, Tokushima, Japan), a food processor for kitchen use (Mini chopper FC-200; Yamazen Co., Osaka, Japan), an ammeter (Clamp on Leak Hitester type 3283; Hioki E.E. Co., Nagano, Japan), a voltage data logger (LabJack U3-HV; Sumitomo Precision Products Co., Ltd., Amagasaki, Japan), and a personal computer with data logging software (DAQ-FactoryExpress, Windows 7/32 bit; Sumitomo Precision Products Co., Ltd., Amagasaki, Japan). The electric power of the food processor was supplied by an electric power source at 50 Hz, and the voltage was limited by the power transformer to control the rotating speed. The torque of the food processor was detected by an ammeter, and the data were captured using a personal computer with a voltage data logger and software. The rotating speed was controlled at 1,000 rpm when running on idle just before sample measurements. Electric current values when running on idle just before sample measurements were subtracted from the data. Data were captured at 0.01-s intervals and smoothed by averaging ± 0.05 s for each mixing time point. Sample measurements were performed three times, and the data were averaged at each mixing time point.

### Mixing-recording measurement of standard fluid

Two types of Newtonian fluids (glycerin and silicone oil) were used as standards. Glycerin viscosity was adjusted by water dilution and temperature changes. Previously reported values (Segur and Oberstar, 1951) were used for the glycerin viscosity at each concentration and temperature. Silicone oil with low viscosity (KF-98-50CS) and high viscosity (KF-98-3000CS) were mixed in various proportions to form a range of viscosities, which were measured with a rheometer (MCR302; Anton-Paar Japan, Tokyo, Japan) having a 25-mm corn plate at a shear rate of 10/s. One hundred milliliters of the fluid was applied to the mixing-recording system, and electric current values were obtained by averaging values from mixing time points at 9-10 s.

### Mixing-recording measurement of agar and gelatin

Samples were dissolved as described for texture measurement and gelled in a plastic container (120 × 90 × 30 mm) overnight at 10°C. Gels were cut into an appropriate size (20-mm cubes) just before measurement, and 100-g samples were placed in the food processor. Paste samples of different sizes were prepared by fracturing sample gels for 120 s using the food processor, and the temperature was readjusted to 10°C. Mixing-recording measurements were performed for 120 s.

### Statistical analysis

Statistical analysis was performed using IBM SPSS (ver. 22) software. Analysis of variance was carried out, followed by Tukey’s post-hoc test.

## RESULTS

### Mixing-recording measurement of standard fluid

The response of the mixing-recording system was investigated using glycerin and silicone oil (Fig. 1). A linear relationship with a high correlation coefficient (r = 0.99) was observed between the electric current values and viscosities.

**FIG. 1.**
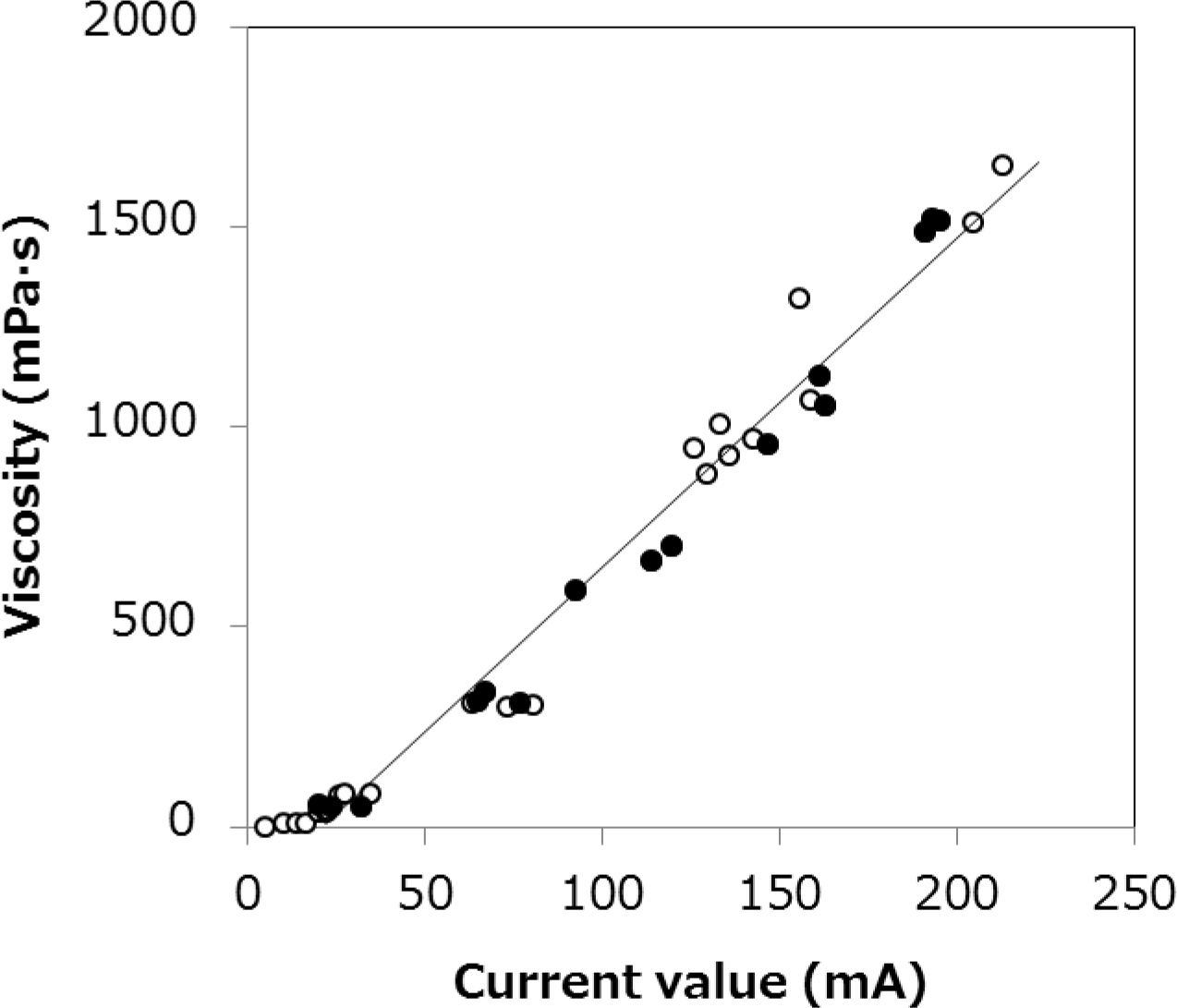
RELATIONSHIP BETWEEN CURRENT VALUE AND VISCOSITY. ○,glycerin; •, silicone oil. Curve-fitting was performed using all data.

### Texture measurement

The texture properties of agar and gelatin at various concentrations are shown in Fig. 2. Agar gels were broken at a strain of 67%, whereas gelatin gels were not. Both gels were broken at a strain of 90%. Adhesiveness values of high-concentration (> 1.0%) agar were high and difficult to evaluate because the gel fragments were attached to the limb of the plunger.

**FIG. 2.**
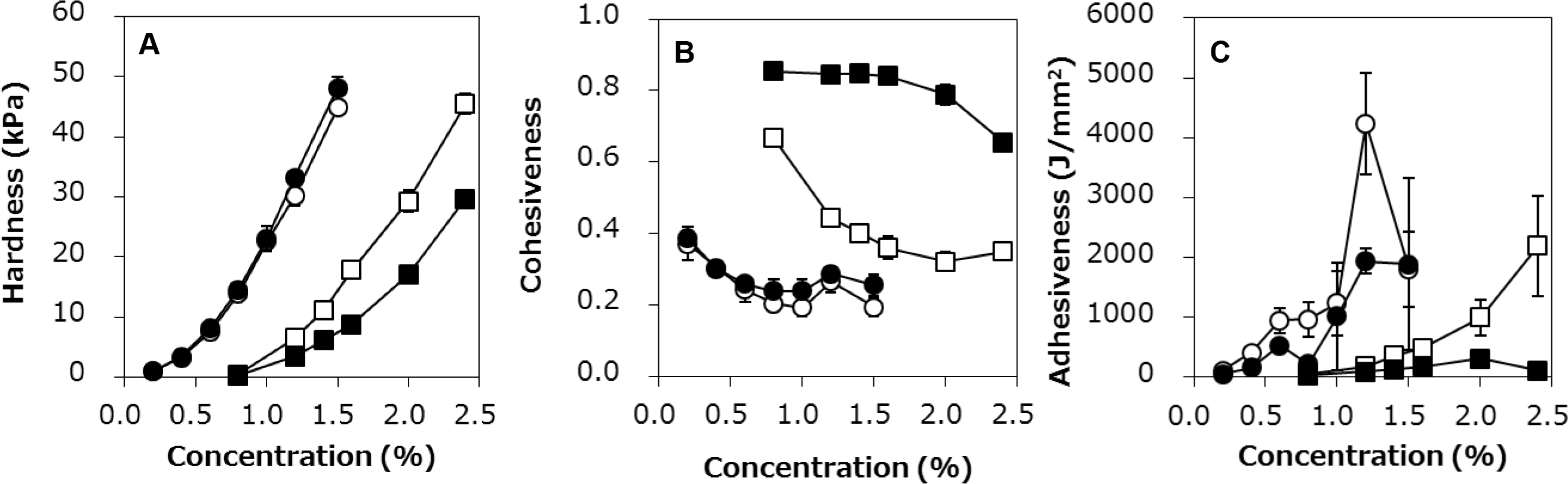
TEXTURE PROPERTIES OF AGAR AND GELATIN. •, agar, 67% strain; ○, agar, 90% strain; ▄, gelatin, 67 % strain; □, gelatin, 90 % strain.

### Mixing-recording measurement of sample gels of various concentrations

Most gel samples were fractured during mixing by the food processor and became smooth pastes, flowing with rotation of the edges. High-concentration (≥ 1.5%) agar did not flow after fracturing. Identical mixing curves for each sample were obtained by monitoring the current value during 120 s of mixing (Fig. 3). The current values of both agar and gelatin decreased during mixing. A sharp decrease was observed in the curves of high-concentration (≥ 1.5%) agar because of racing of the mixer. Fig. 4 shows the current values potted against the concentrations of the gels. The current values increased as the concentration increased, with the exception of those of high-concentration agar at 30 s, which decreased. The high-concentration gelatin values were higher than the agar values at both after 1 and 30 s of mixing.

To investigate the relationship between texture and mixing properties, the current values at certain time points were plotted against the hardness values obtained from TPA at a strain of 90% (Fig. 5). At the beginning of the mixing (time point: 1 s), the current values of gelatin were higher than those of agar (Fig. 5A). In the middle stage of the mixing (time point: 30 s), most sample gels were fractured into pastes. When the current values of gelatin at 30 s were plotted against the hardness values, a linear correlation was observed (Fig. 5B). For low-concentration (≤ 1.0%) agar, the current values were correlated with the hardness values, whereas those of high-concentration (≥ 1.2%) agar were not. Moreover, 1.5% agar did not flow at 30 s, and the current decreased by mixer racing, as described above, although the hardness value was smaller than that of 2.4% gelatin. Data for both gelatin and low-concentration agar were aligned on the same correlation curve with a high correlation coefficient (r = 1.00). The same tendency was observed using the values at 60 and 120 s, with a slightly smaller correlation coefficient (data not shown). No significant relationships were observed between the current values of 30-s mixing and the cohesiveness and adhesiveness values (data not shown).

**FIG. 3.**
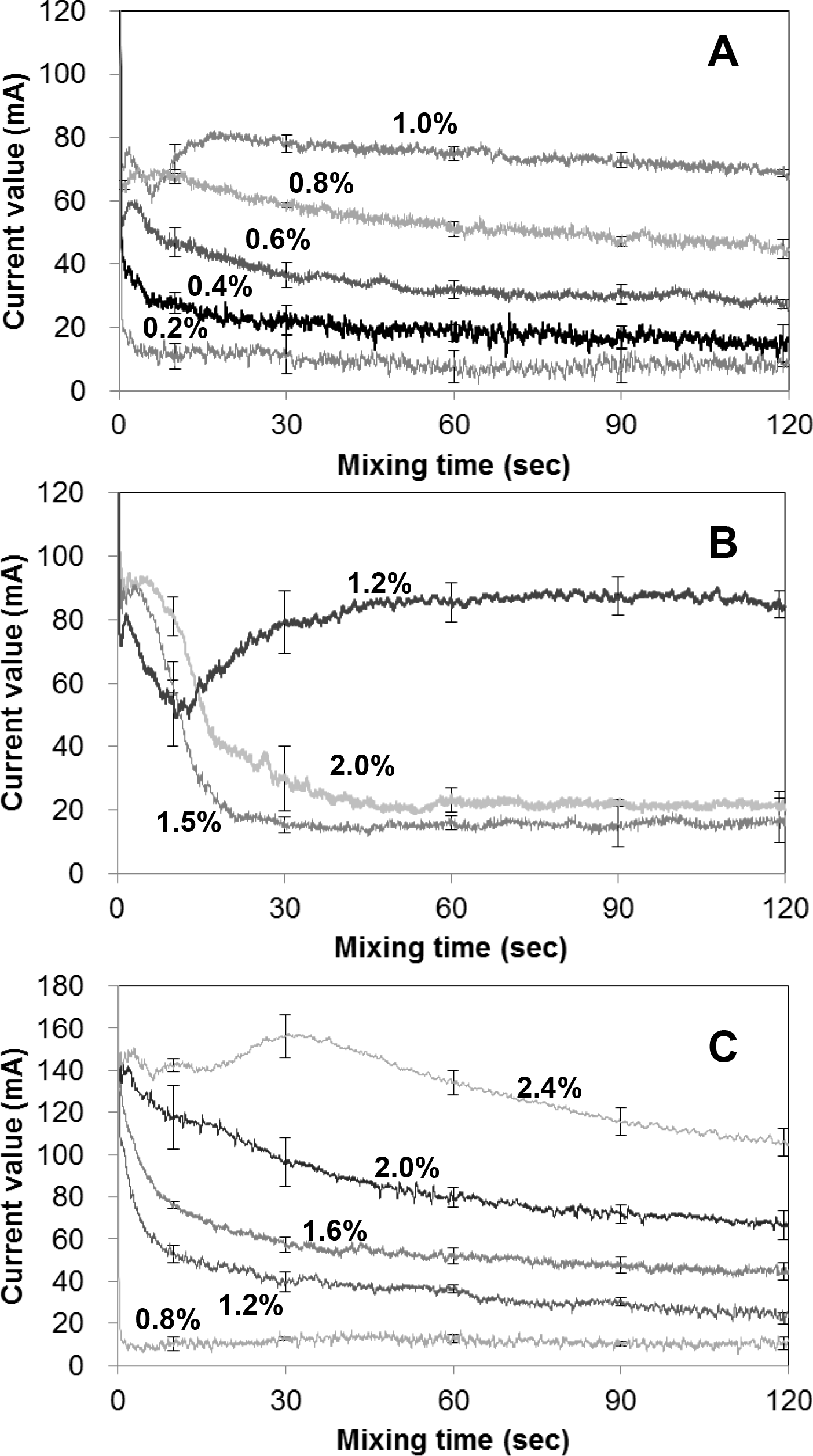
MIXING CURVES OF AGAR AND GELATIN AT DIFFERENT CONCENTRATIONS. A: Agar with a relatively low concentration. B: Agar with a relatively high concentration. C: Gelatin. Error bars represent means ± standard deviations for triplicate experiments averaged for intervals of ± 0.5 s (101 data points) at each mixing time.

**FIG. 4.**
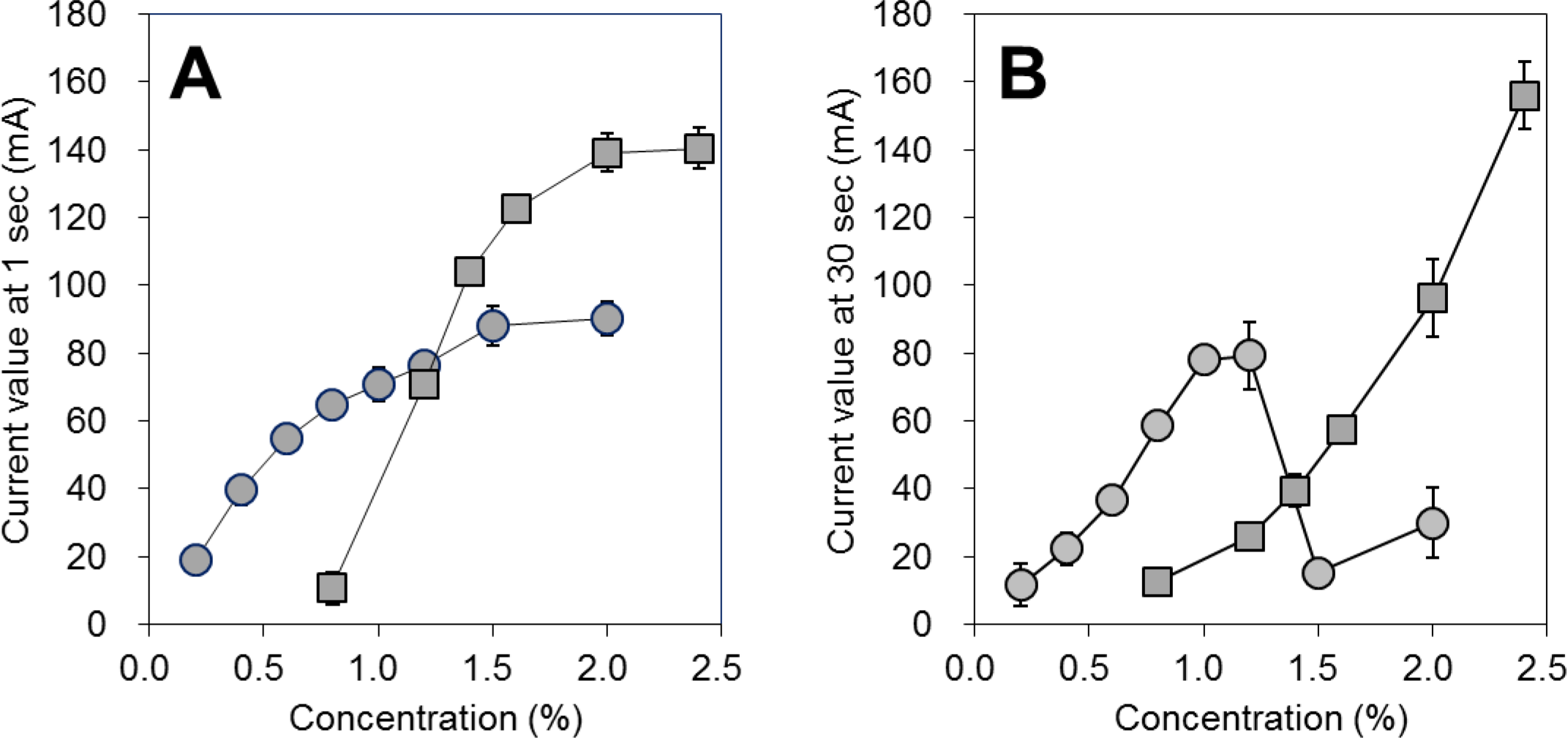
CURRENT VALUES OF AGAR AND GELATIN. •, agar; ▄, gelatin.

**FIG. 5.**
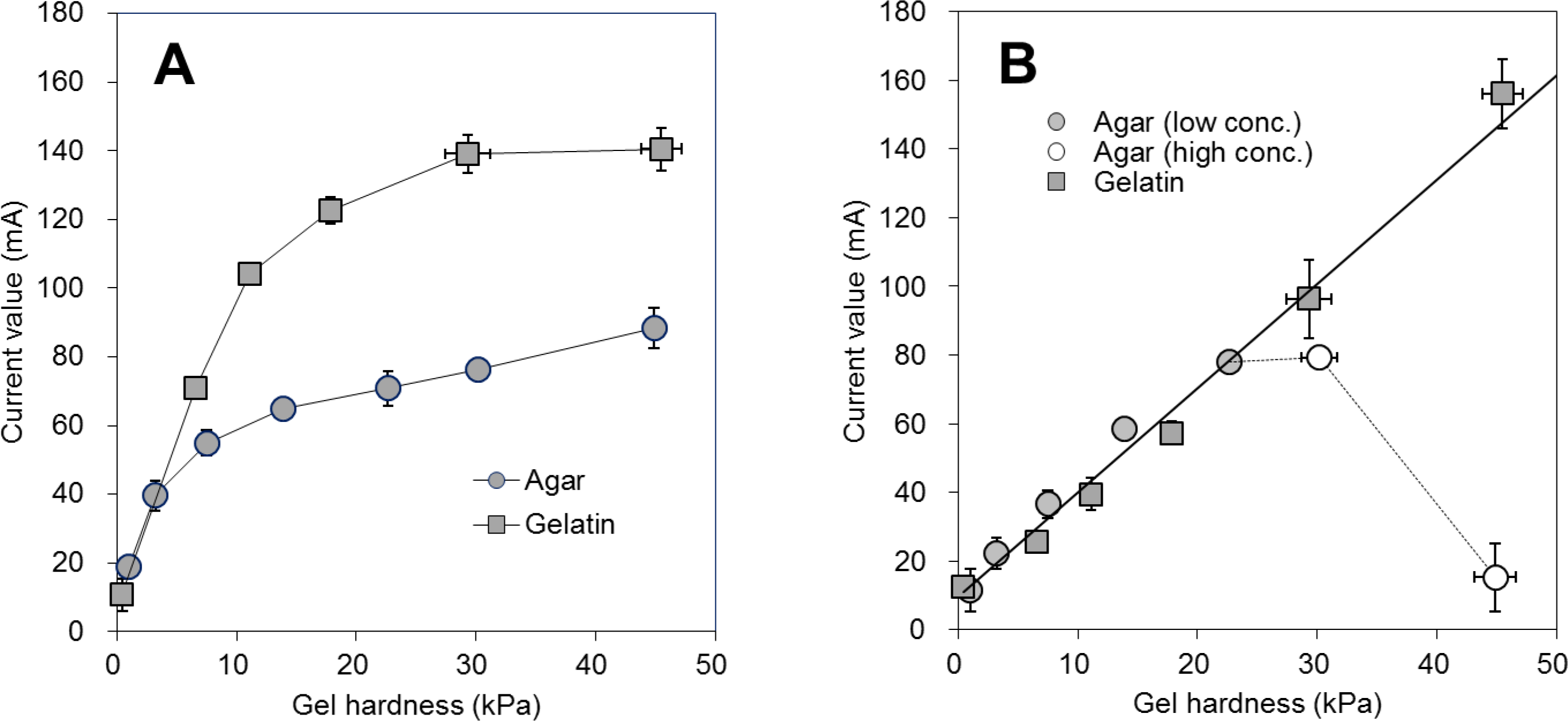
RELATIONSHIP BETWEEN THE CURRENT VALUE OF THE MIXER AND THE GEL HARDNESS VALUE MEASURED BY TPA. A: Current value at 1 s of mixing and gel hardness. B: Current value at 30 s of mixing and gel hardness. Gel hardness was measured at 90% strain. The correlation in B was calculated using agar and gelatin together, except for high-concentration agar.

### Mixing-recording measurement of sample gels of various sizes

Size dependence was investigated using 0.8% agar with 10-mm cubes, 20-mm cubes, and pastes (Fig. 6). Individual mixing curves were obtained, even though the difference between each sample was smaller than that in the former experiment. At the early stage of mixing, the current value increase as the size increased. The difference became smaller during mixing. The differences of integrated current values at each 10-s interval were significant between all samples below 20 s time point and between paste samples and others from 20 to 40 s (*p* < 0.05). No significant differences were observed for all samples at over 40 s of mixing.

**FIG 6.**
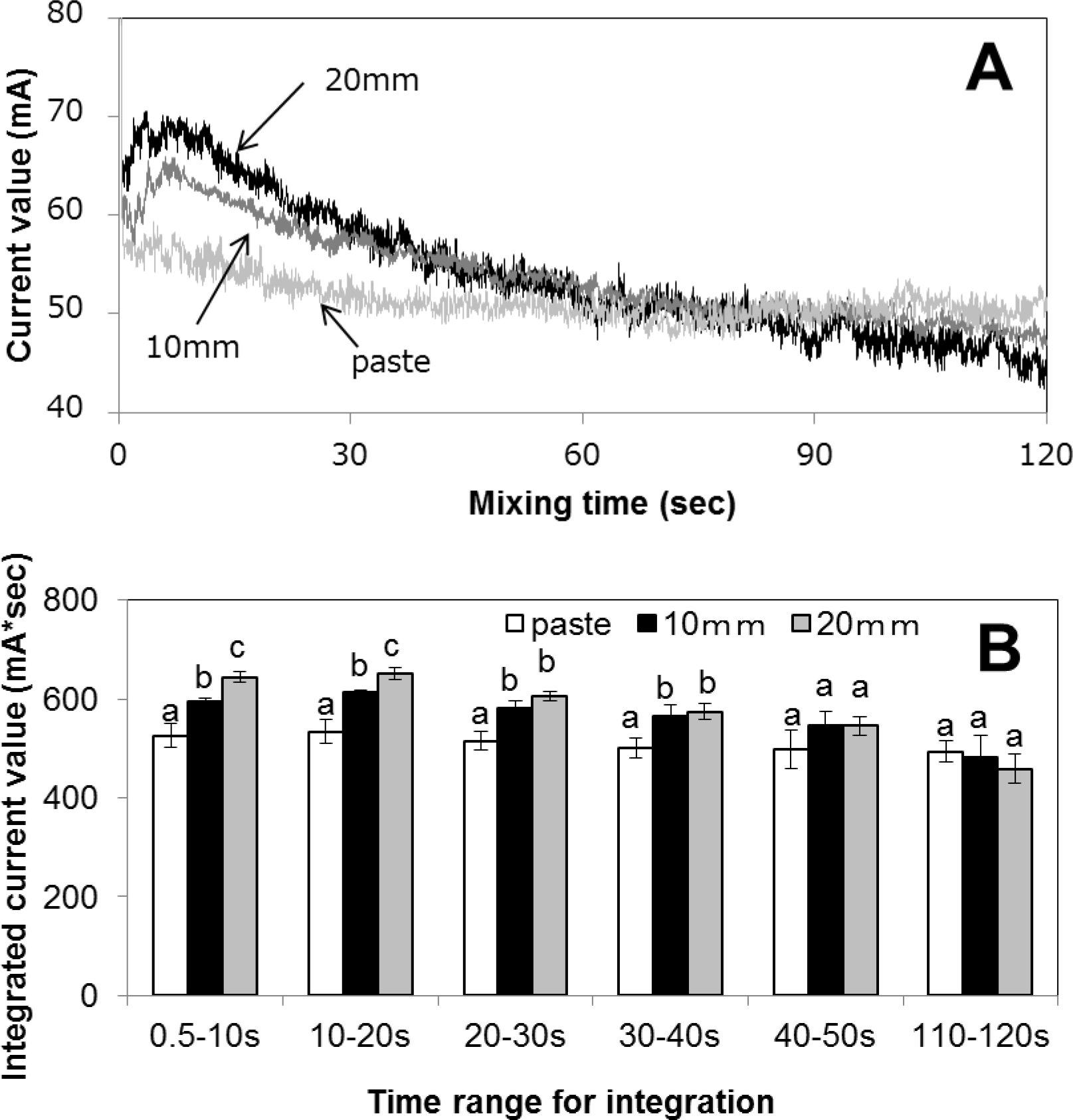
MIXING PROPERTIES OF AGAR GELS OF DIFFERENT SIZES. A: Mixing curves. B: Integrated current values of the mixing curves. Current values were integrated every 10 s. The beginning overcurrent from 0 to 0.5 s was discarded. Error bars represent ± standard deviations for triplicate experiments. Different letters indicate significant difference at *p* < 0.05 in each time range.

## DISCUSSION

In this study, typical Newtonian fluids (glycerin and silicone oil) were applied to a mixing-recording system, and linear regression was observed as described previously (Kanamori, 2016). The current value is thought to indicate some type of viscosity, and the viscosity of liquid foods affects swallowing (Groher, 1987). Because solid foods are swallowed as a bolus, the bolus viscosity is an essential parameter. Thus, mixing-recording measurements of food samples can reveal the viscosity of a mimic bolus of the sample directly and may be suitable for estimating the swallowing properties of food in a simple, reliable manner.

Here, the mixing-recording method was applied to agar and gelatin gels commonly used as gelling agents to study foods for individuals with dysphagia. The results of these samples for TPA at a strain of 90% were similar to those of a previous study (Moritaka and Nakazawa, 2010). Identical mixing curves were obtained from individual gelling agents and concentrations. Solid food is masticated and swallowed as a bolus. The bolus viscosity is affected by the state of fracturing during mastication, which can differ among individuals. Therefore, it is important to measure viscosity at various fracturing rates. The mixing-recording measurement is suitable for this purpose because the viscosities of various fractured states are detected continuously as current values during mixing.

The mixing-recording measurement was time-dependent. In the early stage of mixing, when lumps of gel remained, the current values of gelatin were higher than those of agar. The current value indicated the energy of fracturing gels and may be derived from the physical properties of gels, which was difficult to explain by TPA measurement. In the middle stage of the mixing, the current of 1.5% agar decreased by mixer racing. This could be explained by the observation that the cohesiveness of agar was lower than that of gelatin, even though the cohesiveness values of agar measured by TPA were approximately the same at agar concentrations of 0.6% and 1.5%. Additionally, the low-flow phenomenon appeared in the mixing of relatively hard gels but not in relatively soft gels. The mixing-recording method was able to detect flow or low-flow properties, whereas such measurements were difficult with TPA.

The swallowing properties of gel foods can be estimated by measurement of flow velocity by the pulse Doppler method, as shown previously, and flow velocity has been shown to be correlated with hardness but not cohesiveness or adhesiveness (Moritaka and Nakazawa, 2010; Tanigome et al., 2013). These results were similar to those of the present study, i.e., the current values of relatively soft gels were correlated with the hardness values. It is possible that the bolus viscosity of a soft gel may correspond to the gel hardness. However, in previous studies, experiments were performed with relatively soft gels, as is suitable for foods to be consumed by elderly individuals. The difference between agar and gelatin was clear when harder gels were investigated in the present study. Notably, only a few types of gelling agents have been investigated; therefore, more studies of various types of solid samples are needed.

In this study, larger sample sizes were associated with higher current values during the early stages of mixing, which was reasonable because the energy for mastication is higher as the sample size increases. As the current value of agar decreased during mixing, size reduction before mixing resulted in lower current values at the beginning of mixing and reached a steady-state value earlier. Although food size is one of the most important factors in human mastication and swallowing, conventional methods, such as TPA, are applicable only to samples of an appropriate size. Thus, the mixing-recording method may be useful from the perspective of size-dependent measurement.

Another advantage of the mixing-recording method is simplicity; it is not necessary to prepare materials with different sample sizes and shapes. Actual foods having various sizes and formulas can be applied for the measurement, and the effects of the different sizes and formulas can also provide valuable information. This method could also be applied for food quality control at hospitals and factories, with a short measurement time.

In the present study, a mixing-recording method for the measurement of food texture was established. A commercial food processor was tested as a mixer; however, the equipment used in this study was not optimized. Because food processors function as food fracturing machines, they are suitable for the measurement of actual foods having a wide range of textures. However, other types of mixers, such as pins, paddles, and Z-blades, should be examined for application with different types of samples in future studies.

## CONCLUSION

In this study, a mixing-recording method using a commercial food processor was developed for texture measurement. Identical mixing curves were obtained from the measurement of agar and gelatin gels of several concentrations and sizes. The method was able to estimate the time and size dependence of fracturing, along with various texture and bolus properties. Moreover, this method was also useful for the measurement of food texture, particularly as it related to mastication and swallowing.

## ETHICAL STATEMENTS

Conflict of Interest: The author declares that he has no conflict of interest.

Ethical Review: This study did not involve any human or animal testing.

## ACKNOWLEDGMENTS

The author would like to thank Dr. K. Tsumura (Fuji Oil Holdings Inc.) who offered continuing support and constant encouragement. The author would also like to thank Prof. Jun Kayashita (Prefectural University of Hiroshima) for critical reading of the manuscript.

## REFERENCES

Bloksma, A. H. and Bushuk, W. 1988. Rheology and chemistry of dough. in Wheat Chemistry and Technology, third ed., volume II. (Y. Pomeranz, ed.) pp. 131–217, American Association of Cereal Chemists, St. Paul, America.

Chen, J., 2009. Food oral processing—a review. Food Hydrocolloid., 23, 1–25. doi:10.1016/j.foodhyd.2007.11.013

Friedman, H. H., Whitney, J. E., and Szczesniak, A. S., 1963. The texturometer—a new instrument for objective texture measurement. J. Food Sci., 28, 390–396. doi:10.1111/j.1365-2621.1963.tb00216.x

Fujitani, J., Uyama, R., Okoshi, H., Kayashita, J., Koshiro, A., Takahashi, K., Maeda, H., Fujishima, I. and Ueda, K., 2013. Japanese Society of Dysphagia Rehabilitation: classification of dysphagia modified food. Jpn J Dysphagia Rehabil. 17, 255–67. (in Japanese)

Groher, M. E., 1987. Bolus management and aspiration pneumonia in patients with pseudobulbar dysphagia. Dysphagia, 1, 215–216. doi:10.1007/BF02406920

Hasegawa-Tanigome, A., Ogura, K., Akima, A.,Kohyama, K., Kumagai, H. and Kumagai, H., 2013. Relationship between the parameters obtained by 2-bite texture profile analysis (TPA) and the velocity through the pharynx measured by the ultrasonic pulse Doppler method. Japan J. Food Eng., 14, 87–96. doi:10.11301/jsfe.14.87(in Japanese)

Kanamori, J., 2016. Cereal Chem., in press. doi:10.1094/CCHEM-12-15-0258-R

Kumagai, H. and Tanigome, A., 2011. Enge syogaisya you kaigosyoku no tekusutyah · bussei (Texture and physical properties of food for dysphagia.). in Sinka suru syokuhin tekusutyah kenkyuu (Developping studies of food texture.) (Yamano, Y eds.) NTS, Tokyo, Japan. (in Japanese)

Michiwaki, Y., Yokoyama, M., Michi, K., Ohkoshi, H., Takahashi, T. and Hirota, E., 2000. Texture analysis of diets for patients with dysphagia. Jpn. J. Dysphagia Rehabil., 4, 28–32. (in Japanese)

MINISTRY OF HEALTH, LABOUR AND WELFARE, 2009. Indication Permission of Food Safety Bureau and Department of Food Safety. No.051101 (placed under Consumer Affairs Agency, Government of Japan from 2010) (in Japanese)

Moritaka, H., and Nakazawa, F., 2010. Flow velocity of a bolus in the pharynx and rheological properties of agar and gelatin. J. Texture Stud., 41, 139–152. doi:10.1111/j.1745-4603.2010.00218.x19

NATIONAL DYSPHAGIA DIET TASK FORCE, AND AMERICAN DIETETIC ASSOCIATION, 2002. National Dysphagia Diet: Standardization for Optimal Care. American Dietetic Associati.

Nishinari, K., Kohyama, K., Kumagai, H., Funami, T., and Bourne, M. C. 2013. Parameters of texture profile analysis. Food Sci. Tech. Res., 19, 519–521. doi:10.3136/fstr. 19.519

Sakai, M., Egashira, F., Kanaya, S. and Kayashita, J., 2006. Comparison of food physical properties with reference to clinically effective stepwise swallowing foods. Jpn. J. Dysphagia Rehabil., 10, 239–248. (in Japanese)

Segur, J. B., and Oberstar, H. E., 1951. Viscosity of glycerol and its aqueous solutions. Ind. Eng. Chem., 43, 2117–2120. doi:10.1021/ie50501a040

Yamagata, Y. and Kayashita, J., Evaluation of the Japanese Dysphasia Diet 2013 by the JSDR Dysphagia Diet Committee (thickened liquid) by using several types of thickened liquids. Jpn. J. Dysphagia Rehabil., 19, 109–116. (in Japanese)

